# Learning the synaptic and intrinsic membrane dynamics underlying working memory in spiking neural network models

**DOI:** 10.1101/2020.06.11.147405

**Authors:** Yinghao Li, Robert Kim, Terrence J. Sejnowski

## Abstract

Recurrent neural network (RNN) model trained to perform cognitive tasks is a useful computational tool for understanding how cortical circuits execute complex computations. However, these models are often composed of units that interact with one another using continuous signals and overlook parameters intrinsic to spiking neurons. Here, we developed a method to directly train not only synaptic-related variables but also membrane-related parameters of a spiking RNN model. Training our model on a wide range of cognitive tasks resulted in diverse yet task-specific synaptic and membrane parameters. We also show that fast membrane time constants and slow synaptic decay dynamics naturally emerge from our model when it is trained on tasks associated with working memory (WM). Further dissecting the optimized parameters revealed that fast membrane properties and slow synaptic dynamics are important for encoding stimuli and WM maintenance, respectively. This approach offers a unique window into how connectivity patterns and intrinsic neuronal properties contribute to complex dynamics in neural populations.

## Introduction

Neurons in the cortex form recurrent connections that give rise to the complex dynamic processes underlying computational functions [1–4]. Previous studies have used models based on recurrent neural networks (RNNs) of continuous-rate units to characterize network dynamics behind neural computations and to validate experimental findings [5–10]. However, these models do not explain how intrinsic membrane properties could also contribute to the emerging dynamics.

Rate-based encoding of information has been reliably observed in experimental settings [8]. However, recent studies demonstrated that membrane potential dynamics along with spike-based coding are also capable of reliably transmitting information [11–13]. In addition, the intrinsic membrane properties of inhibitory neurons, including the membrane time constant and rheobase (minimum current required to evoke a single action potential), were different in two higher-order cortical areas [14]. These findings strongly indicate that neuronal intrinsic properties, often ignored in previous computational studies employing rate-based RNNs, are crucial for better understanding how distinct subtypes of neurons contribute to information processing.

Rate-based RNNs can be easily trained by stochastic gradient-descent to perform specified cognitive tasks [15]. However, similar supervised learning methods cannot be used to train spiking RNNs due to the non-differentiable behavior of action potentials [16]. Thus, several methods introduced differentiable approximations of the non-differentiable spiking dynamics [17–20]. These studies directly applied backpropagation to tune synaptic connections for task-specific computations. Other methods that do not rely on gradient computations have been also utilized to train spiking networks. One such method is based on the first-order reduced and controlled error (FORCE) algorithm previously developed for rate RNNs [6]. The FORCE-based methods are capable of training spiking networks, but training all the parameters including recurrent connections could become computationally inefficient [21–23]. Lastly, recent studies successfully converted rate-based networks trained with a gradient-descent method to spiking networks for both convolutional and recurrent neural networks [24, 25]. Since these models are built on rate-coding networks, the resulting spiking models do not take advantage of the rich spiking dynamics. Moreover, these previous models assume that all the units in a trained network are equivalent, even though experimental evidence shows that neurons in biological neural networks are highly heterogeneous. Such diversity has a vital role in efficient neural coding [26].

Here, we present a new approach that can directly train not only recurrent synapses but also membrane-related parameters of a spiking RNN model. Our method utilizes mollifier functions [27] to alter the spiking dynamics to be differentiable, and a gradient-descent method is applied to tune the model parameters. These parameters are composed of synaptic parameters including recurrent connections and several important spiking-related parameters such as membrane time constant and action potential threshold. Neurons with diverse and heterogeneous intrinsic parameters emerged from training our spiking model on a wide range of cognitive tasks. Furthermore, we observed that both synaptic and spiking parameters worked in a synergistic manner to perform complex tasks that required information integration and working memory.

## Results

Here, we provide an overview of the method that we developed to directly train spiking recurrent neural network (RNN) models (for more details see Methods). Throughout the study, we considered recurrent network models composed of leaky integrate-and-fire (LIF) units whose membrane voltage dynamics were governed by:

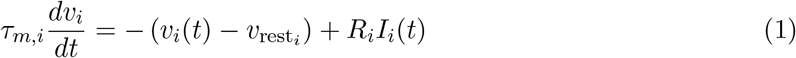

where *τ*_*m,i*_ is the membrane time constant of unit *i*, *v*_*i*_(*t*) is the membrane voltage of unit *i* at time *t*, *v*_*rest,i*_ is the resting potential of unit *i*, and *R*_*i*_ is the input resistance of unit *i*. *I*_*i*_(*t*) represents the current input to unit *i* at time *t*, which is given by:

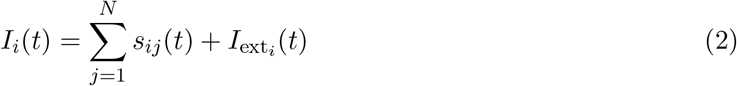

where *N* is the total number of units in the network, *s*_*ij*_(*t*) is the synaptic input from unit *j* to unit *i* at time *t*, and *I*_*ext,i*_(*t*) is the external current source into unit *i* at time *t*. We used a single exponential synaptic filter to model the synaptic input (*s*):

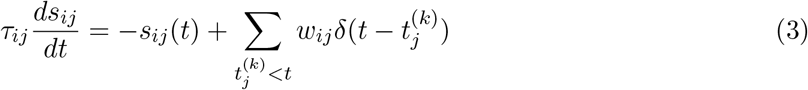

where *τ*_*ij*_ is the decay time constant of the synaptic current from unit *j* to unit *i*, *w*_*ij*_ is the synaptic strength from unit *j* to unit *i*, 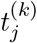 denotes the time of the *k*-th action potential of unit *j*, and *δ*(*x*) is the Dirac delta function. Once the membrane voltage of the unit *i* crosses its action potential threshold (*ϑ*_*i*_), its membrane voltage is brought back down to its reset voltage (*v*_*reset,i*_).

Each LIF unit is characterized by five distinct parameters: membrane time constant (*τ*_*m,i*_), resting potential (*v*_*rest,i*_), input resistance (*R*_*i*_), action potential threshold (*ϑ*_*i*_), and reset potential (*v*_*reset,i*_). In addition, there are two trainable synaptic parameters: synaptic strength (*w*_*ij*_) and synaptic decay time constant (*τ*_*ij*_) from unit *j* to unit *i*.

In order to tune all the parameters described above to produce functional spiking RNNs capable of performing cognitive tasks, we employed the commonly used gradient-descent method known as backpropagation through time (BPTT; [28]) with a few important modifications. We utilized mollifier gradient approximations to avoid the non-differentiability problem associated with training spiking networks with backpropagation [27]. Furthermore, we optimized each of the model parameters (except for the synaptic connectivity weights) in a biologically plausible range (see Methods). We also employed the weight parametrization method proposed by Song et al. to impose Dale’s principle [29] (see Methods). All the spiking RNN models trained in the study used the parameter value ranges listed in Supplementary Table 1 unless otherwise noted.

### Units with diverse parameter values emerge after training

We applied our method to train spiking networks to perform the context-dependent input integration task previously employed by Mante et al. [8]. Briefly, Mante et al. trained rhesus monkeys to flexibly integrate sensory inputs (color and motion of randomly moving dots presented on a screen). A contextual cue was given to instruct the monkeys which sensory modality (color or motion) they should attend to. The monkeys were required to employ flexible computations as the same modality could be either relevant or irrelevant depending on the contextual cue. Several previous modeling studies have successfully implemented a simplified version of the task and reproduced the neural dynamics present in the experimental data with both continuous-rate RNNs and spiking RNNs converted from rate RNNs [25, 29, 30]. With our method, we were able to directly train the first, to our knowledge, spiking RNNs with heterogeneous units whose parameters were within biologically plausible limits.

In order to train spiking RNNs to perform the input integration task, we employed a task paradigm similar to the one used by previous computational studies [8, 25, 29, 30]. A recurrently connected network received two streams of noisy input signals along with a constant-valued signal that encoded the contextual cue (Fig. 1A). The input signals were sampled from a standard Gaussian distribution (i.e., with zero mean and unit variance) and then shifted by a positive or negative “offset” value to simulate the evidence presented in the input modalities. The network was trained to produce an output signal approaching either +1 or −1 depending on the cue and the evidence present in the input signal: if the cued input had a positive mean, the output signal approached +1, and vice versa (Fig. 1B top). The input signal, 150 ms in duration, was given after a fixation period (300 ms), and the network was trained to produce an output signal immediately after the offset of the input signal.

**Fig. 1.**
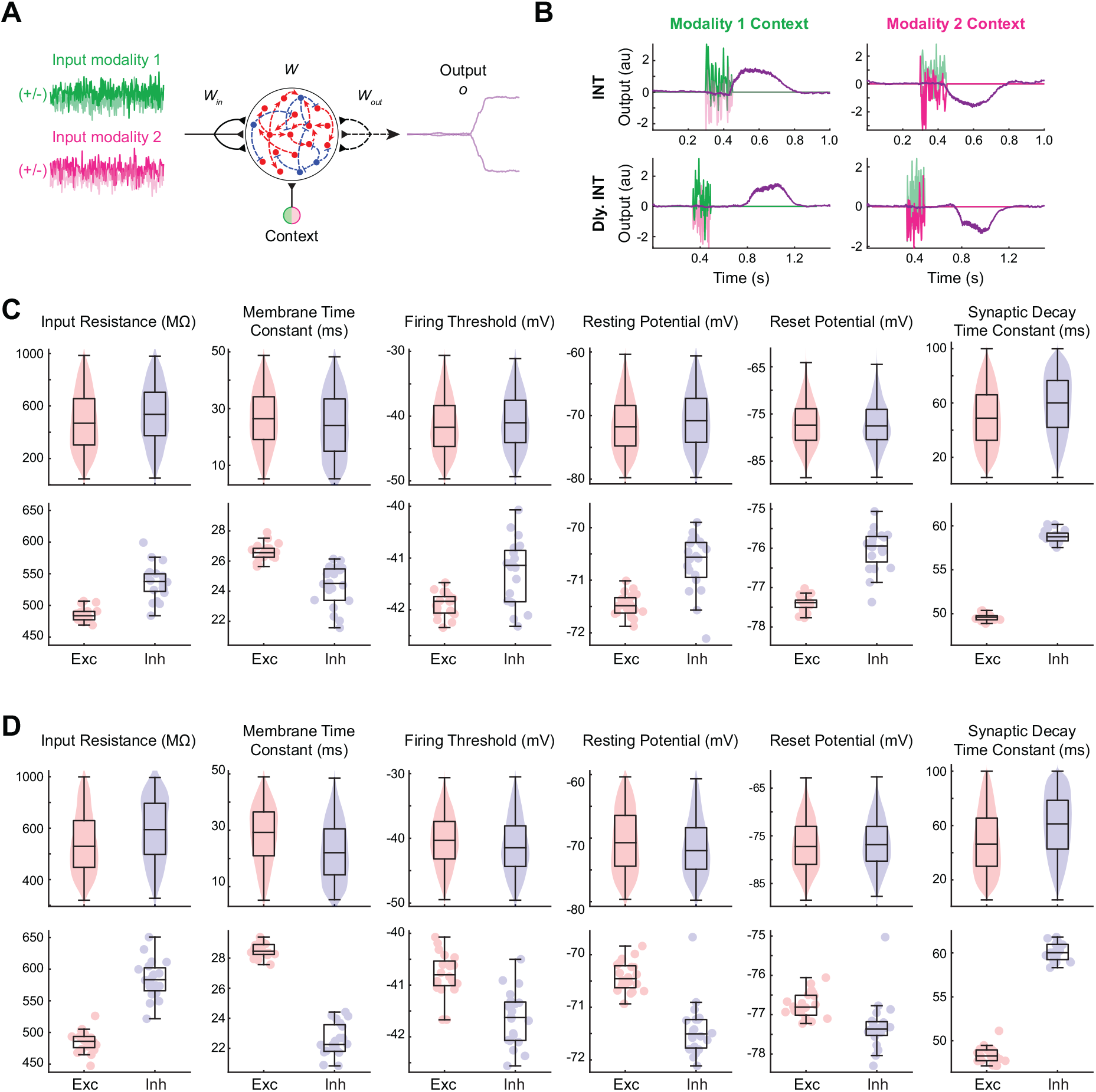
Biologically realistic spiking network performing a context-dependent input integration task. (A) Schematic diagram of the RNN model trained for the context-dependent integration task. Two streams of noisy input signals (green and magenta lines) along with a context signal were delivered to the LIF RNN. The network was trained to integrate and determine if the mean of the cued input signal (i.e., cued offset value) was positive (“+” choice) or negative (“_” choice) without or with a delay period at the end of the noisy input signals. (B) Example input and output signals from example RNNs trained to perform the task without (top row; INT) or with a delay period (bottom row; Dly. INT). (C) Distributions of the optimized parameters for the excitatory (red) and inhibitory (blue) units across all 20 models trained for the INT task. Top, distributions pooled from all the units from 20 models. Bottom, each dot represents the average value from one network. (D) Distributions of the optimized parameters for the excitatory (red) and inhibitory (blue) units across all 20 models trained for the Dly. INT task. Top, distributions pooled from all the units from 20 models. Bottom, each dot represents the average value from one network.

We trained 20 spiking RNNs to perform the context-based input integration task. All the trainable parameters were initialized with random numbers drawn from a standard Gaussian distribution and re-scaled to the biologically plausible ranges (see Methods and Supplementary Table 1). Each network was trained until the training termination criteria were satisfied (see Methods). On average, 508.21 ± 45.96 training trials were needed for a network to meet the training termination conditions. After training, a wide distribution of the parameters emerged for both excitatory and inhibitory populations (Fig. 1C, top).

Consistent with the previous experimental recordings from cortical neurons, the inhibitory units in our trained RNNs fired at a higher rate compared to the excitatory units [31]. The higher average firing rates of the inhibitory units were largely due to the intrinsic properties that resulted from training. Compared to the excitatory population, the inhibitory units in the trained RNNs had significantly larger input resistance, smaller membrane time constants, and more depolarized resting potential (Fig. 1C; *P* < 0.0001, two-sided Wilcoxon rank-sum test). The action potential thresholds and the reset potentials were significantly more depolarized for the inhibitory group. Furthermore, the time constants of the inhibitory synaptic current variable were significantly larger than the excitatory synaptic decay time constants (Fig. 1C).

### Working memory requires distinct parameter distributions

The context-dependent input integration task considered in the previous section did not require complex cognitive skills such as working memory (WM) computations. In order to explore what parameter values are essential for WM tasks, we modified the paradigm to incorporate a WM component by adding a delay period after the delivery of the input signals. The RNN model was trained to integrate the noisy input signals, sustain the integrated information throughout the 300 ms delay period, and produce an output signal (Fig. 1B bottom). We again trained 20 models for the modified integration task with the same training termination criteria (see Methods). This task required more training trials (on average 1618.10 ± 345.54), but all the models were successfully trained within 2000 training trials.

Overall, the distributions of the trained parameters were similar to those observed from the RNNs trained on the non-WM version of the task (Fig. 1D). The parameters that were significantly different between the two RNN models were the membrane time constant and the synaptic decay time constant. The inhibitory units from the WM model displayed much faster membrane dynamics and slower synaptic decay compared to the inhibitory population of the non-WM model (*P* < 0.0001, two-sided Wilcoxon rank-sum test).

To ensure that the patterns of the trained parameters and the distinct distributions of the two parameters (*τ*_*m*_ and *τ*) observed from the delayed integration model were indeed associated with WM computations, we trained RNNs on two additional WM-related tasks: delayed matched-to-sample (DMS) and delayed discrimination (DIS) tasks. For each task, we again trained 20 RNNs. Both task paradigms included two sequential stimuli separated by a brief delay period. For the DMS task, the two input stimuli were either +1 or −1; if the two sequential had the same sign (i.e., +1*/* + 1 or −1*/* − 1), the network was trained to have an output signal approaching +1, while if the two stimuli had different signs (i.e., +1*/* − 1 or −1*/* + 1), the output signal approached −1 (Fig. 2A; see Methods). The two input stimuli for the DIS task were sinusoidal waves with different frequencies, modeled after the task used by Romo et al. [32] where monkeys were trained to discriminate two vibratory stimuli. If the first stimulus had a higher (lower) frequency, our RNN model was trained to produce a positive (negative) output signal (Fig. 2B; see Methods).

**Fig. 2.**
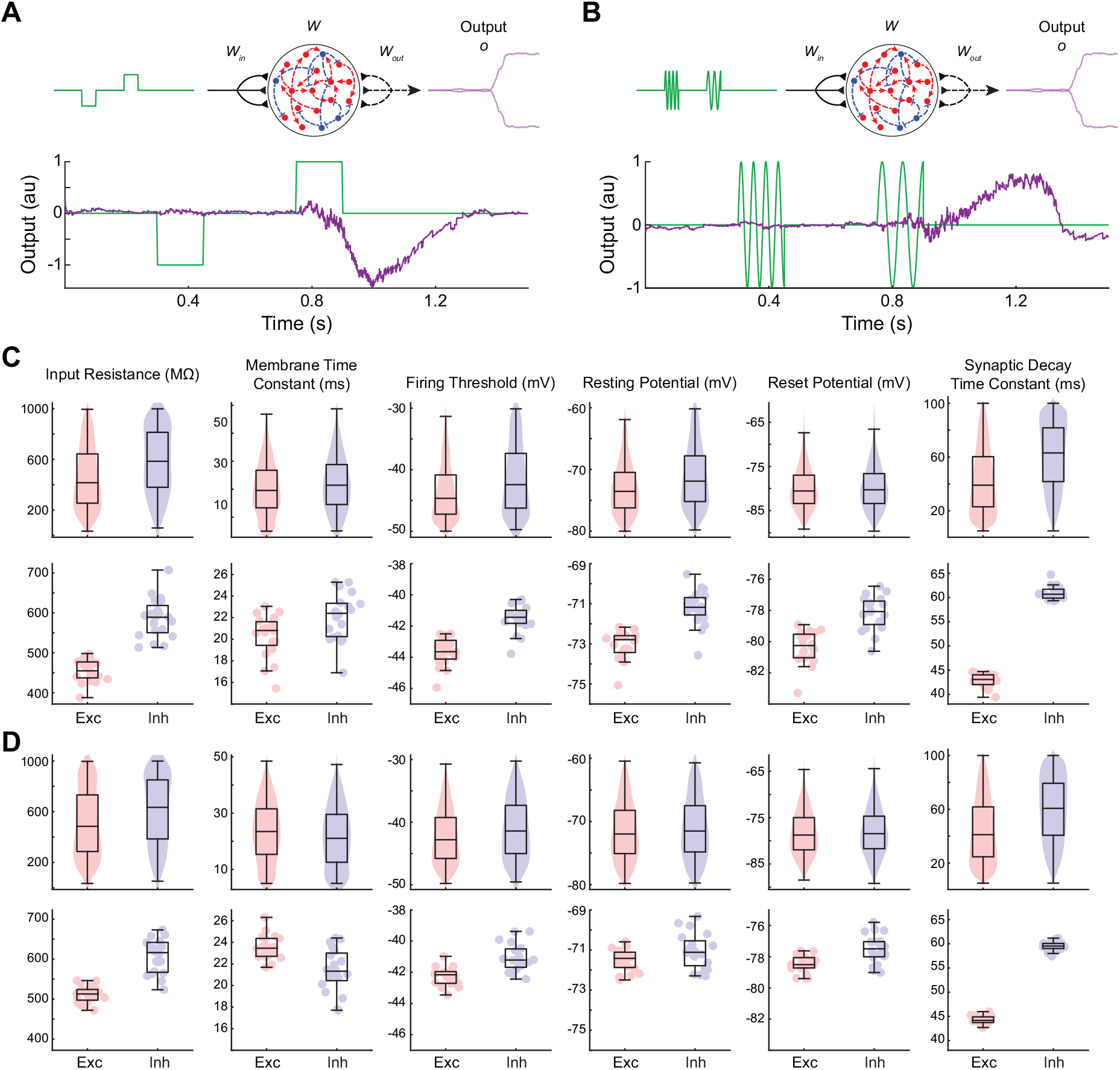
RNNs trained for two additional WM tasks. (A) Schematic illustrating the task paradigm for the delayed match-to-sample (DMS) task (top) and input and output signals from an example trained RNN (bottom). (B) Schematic illustrating the task paradigm for the delayed discrimination (DIS) task (top) and input and output signals from an example trained RNN (bottom). (C) Distributions of the optimized parameters for the excitatory (red) and inhibitory (blue) units across all 20 models trained for the DMS task. Top, distributions pooled from all the units from 20 models. Bottom, each dot represents the average value from one network. (D) Distributions of the optimized parameters for the excitatory (red) and inhibitory (blue) units across all 20 models trained for the DIS task. Top, distributions pooled from all the units from 20 models. Bottom, each dot represents the average value from one network.

It took longer to train our model on these two tasks compared to the delayed integration task (7103.95 ± 3738.65 trials for the DMS task and 6985.47 ± 2112.34 trials for the DIS task). The distributions of the tuned parameters from the two WM tasks were similar to the distributions obtained from the delayed integration task (Fig. 2C and D). More importantly, we again observed significantly faster membrane voltage dynamics and slower synaptic decay from the inhibitory units in the DMS and DIS models compared to the inhibitory units from the non-WM task. These findings strongly suggest that the two parameters (*τ*_*m*_ and *τ*) of the inhibitory group contribute to important dynamics associated with WM.

### Shared intrinsic properties across different working memory tasks

Prefrontal cortex and other higher-order cortical areas have been shown to integrate information in a flexible manner and switch between tasks seamlessly [8]. Along this line of thought, we hypothesized that the intrinsic properties optimized for one WM task should be generalizable to other tasks that also require WM. In order to test this hypothesis, we re-trained all the RNNs that were trained in the previous sections to perform the DMS task without tuning the intrinsic parameters. For example, given a network trained on the non-WM integration task, we froze its intrinsic (*R*, *τ*_*m*_, *v*_rest_, *v*_reset_ *ϑ*) along with the synaptic decay time constant (*τ*) and optimized the recurrent connections (*W*) only using BPTT (see Methods). Therefore, each of the 20 RNNs trained for each of the four tasks (non-WM integration, delayed integration, DMS, and DIS tasks) was re-trained to perform the DMS task. As expected, the average number of trials required to successfully retrain the RNNs previously trained for the DMS task was low at 4408.95 ± 3596.27 (Fig. 3A). The number of trials required to re-train the RNNs from the DIS task was also low at 4180.30 ± 2692.81. The RNNs trained for the delayed integration task took longer to re-train at 5391.85 ± 2197.99. The non-WM RNNs required the most number of training trials to perform the DMS task (9647.55 ± 2933.17). These findings indicate that the intrinsic properties from one WM model are transferable to other WM models.

**Fig. 3.**
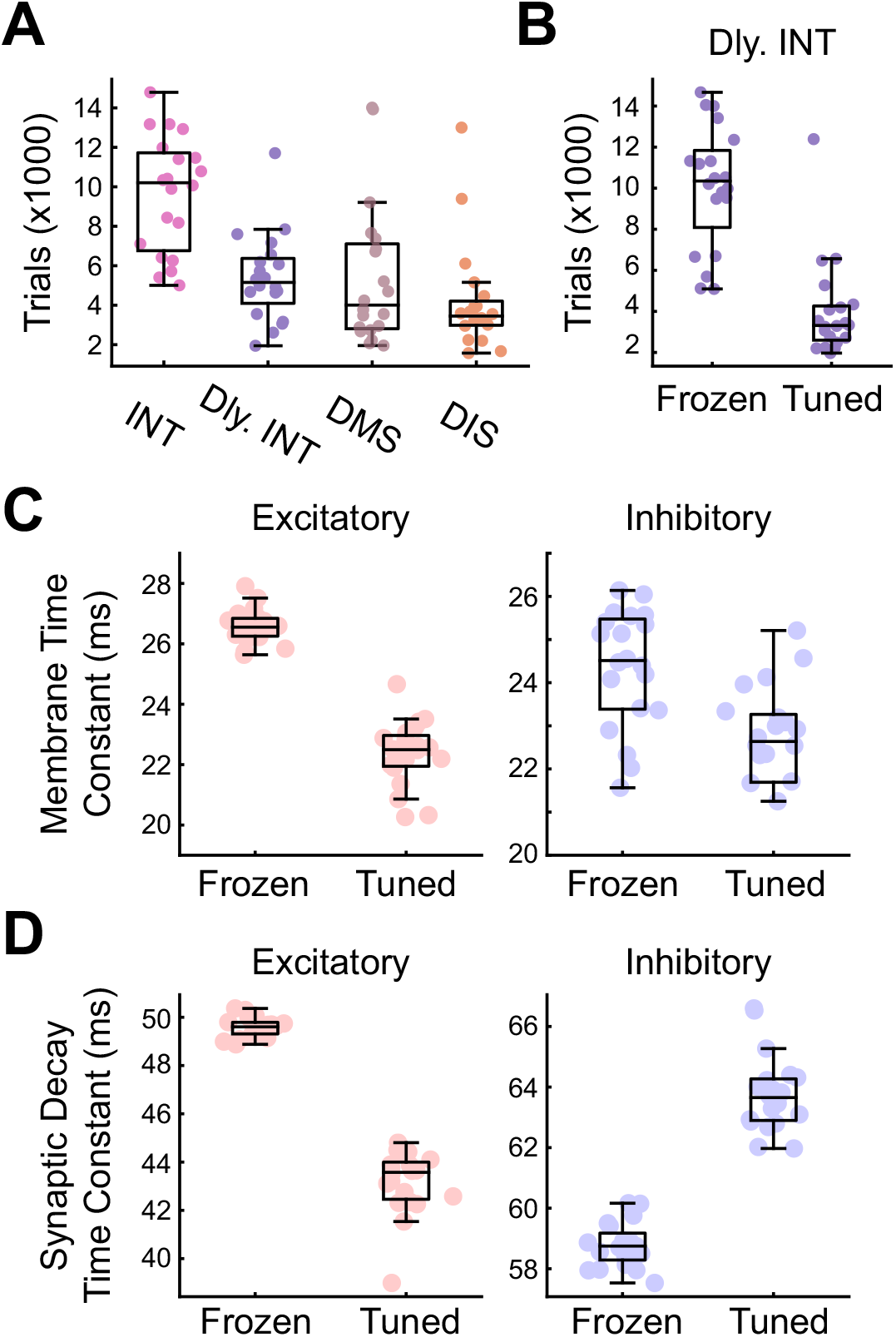
Retraining RNN models to perform the DMS task. (A) Number of training trials required to retrain the models previously trained for the INT, Dly. INT, DMS, or DIS tasks to perform the DMS task. (B) Number of training trials required to retrain the Dly. INT RNNs to perform the DMS task with the membrane time constant (*τ*_*m*_) and synaptic decay time constant (*τ*) frozen or tuned. (C) Distribution of the membrane time constant values for the excitatory (red) and inhibitory (blue) units for the two conditions (frozen and tuned). Each dot represents the average value from one network. (D) Distribution of the synaptic decay time constant values for the excitatory (red) and inhibitory (blue) units for the two conditions (frozen and tuned). Each dot represents the average value from one network.

Based on these previous results, the membrane time constant (*τ*_*m*_) and the synaptic decay (*τ*) variables appeared to be the two most important parameters for the transferability of WM. To test this, we repeated the re-training procedure with both *τ*_*m*_ and *τ* either fixed (“frozen”) or optimized (“tuned”) for the non-WM RNNs (see Methods). For the “frozen” condition (i.e., *τ*_*m*_ and *τ* frozen while the other parameters optimized), the number of trials required to re-train the non-WM RNNs to perform the DMS task was high and not significantly different from the number of trials it took with the intrinsic parameters fixed (Fig. 3B). On the other hand, re-tuning only *τ*_*m*_ and *τ* with the other parameters fixed (i.e., “tuned” condition) resulted in a significant reduction in training time (Fig. 3B), suggesting that these two parameters are indeed critical for performing WM. Optimizing both *τ*_*m*_ and *τ* resulted in a significant decrease in *τ*_*m*_ for both excitatory and inhibitory populations (Fig. 3C). The synaptic decay values decreased for the excitatory units after re-tuning (Fig. 3D left). For the inhibitory population, *τ* was significantly increased (Fig. 3D right).

### Membrane and synaptic decay time constants critical for WM maintenance

Pyramidal excitatory neurons and parvalbumin (PV) interneurons make up the majority of the neuronal cell population in the cortex, and they have been shown to specialize in fast and reliable encoding of information with high temporal precision [33]. To further investigate if the fast membrane and slow synaptic dynamics of the units from our WM RNNs are aligned with previous experimental findings and to probe how they contribute to WM maintenance, we manipulated *τ*_*m*_ and *τ* during different epochs of the DMS task paradigm.

For each of the RNNs trained from the DMS task, we first divided the population into two subgroups based on their *τ*_*m*_ values (see Methods). The short *τ*_*m*_ group contained units whose *τ*_*m*_ was smaller than the lower quartile value, while the long *τ*_*m*_ group contained units whose *τ*_*m*_ was greater than the upper quartile. During each of the four epochs (fixation, first stimulus, delay, and second stimulus), we then inhibited the two *τ*_*m*_ subgroups separately by hyperpolarizing them and assessed the task performance (see Methods). As shown in Fig. 4, inhibiting the short *τ*_*m*_ subgroup during the two stimulus windows significantly impaired task performance (Fig. 4B and D), while disrupting the long *τ*_*m*_ group did not result in significant changes in task performance in all four task epochs.

**Fig. 4.**
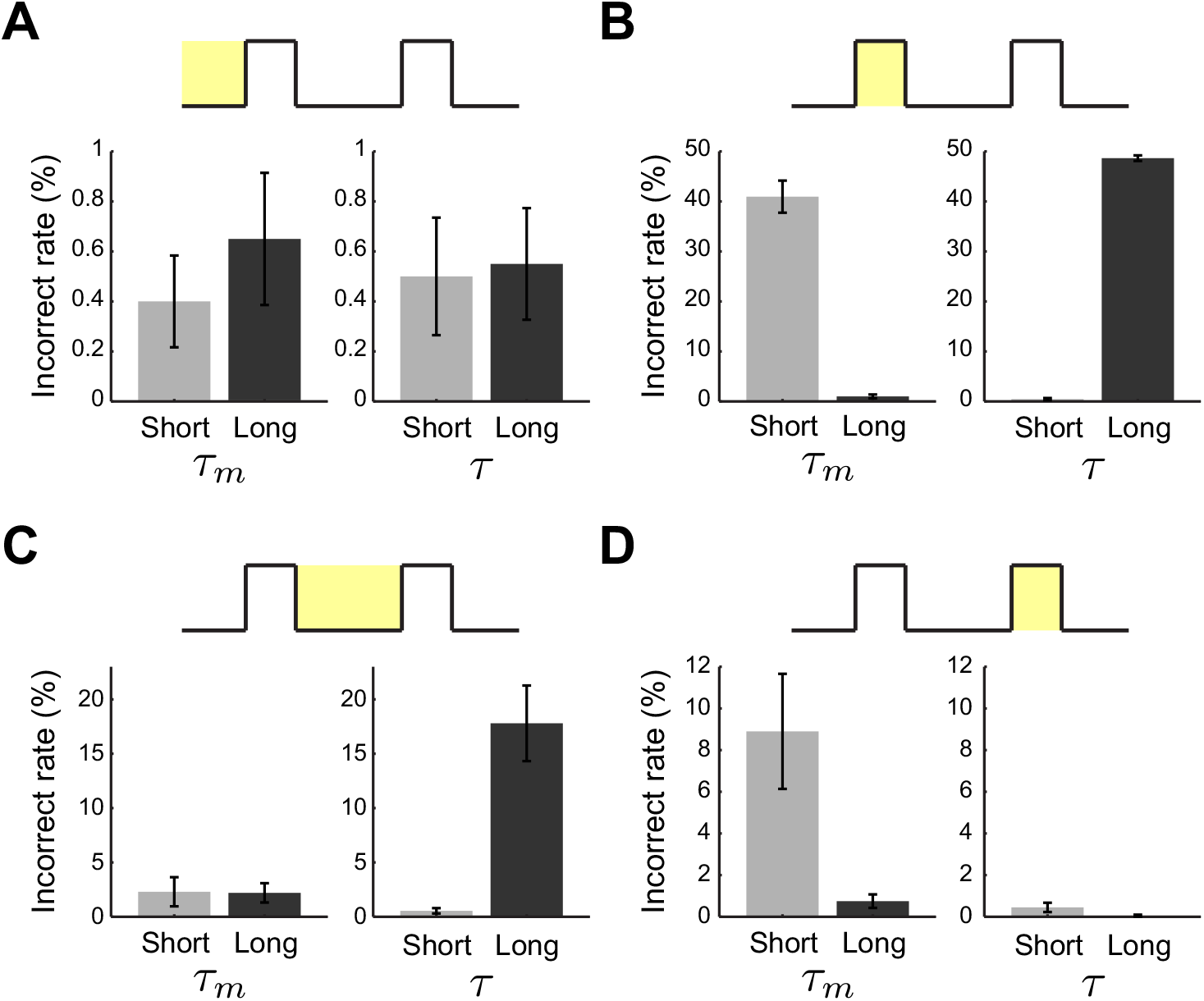
Membrane and synaptic time constants important for encoding stimuli and WM maintenance. (A–D) DMS task performance when short *τ*_*m*_, long *τ*_*m*_, short *τ*, or long *τ* units were inhibited during the fixation (A), first stimulus window (B), delay period (C), or second stimulus window (D).

We repeated the above analysis with two subgroups derived from a quartile split of the synaptic decay time constant (*τ*; see Methods). Suppressing the synaptic connections in the long *τ* subgroup during the first stimulus window and the delay period significantly impaired task performance (Fig. 4B and C). Inhibiting the short *τ* group at any of the four epochs did not affect the task performance.

Therefore, the units with the fast membrane voltage dynamics (*τ*_*m*_) were important for encoding of stimuli, while the slow synaptic dynamics (*τ*) were critical for maintaining the first stimulus information throughout the period spanning from the first stimulus window to the end of the delay window.

## Discussion

In this study, we presented a new method for directly training spiking RNNs with a gradient-based supervised training algorithm. Our approach allows optimizing not only the synaptic variables but also parameters intrinsic to spiking dynamics. By optimizing a wide range of parameters, we first demonstrated that units with diverse features emerged when the model was trained on a cognitive task (Figs. 1 and 2). We also showed that fast membrane dynamics combined with a slow synaptic property are critical for performing WM tasks (Figs. 3 and 4). Diversity is a basic biological principle that emerged here as a basic computational principle in spiking neural models.

Previous modeling studies have trained RNNs to perform cognitive tasks [8, 34, 35]. Although some of these studies were able to train spiking RNN models, the intrinsic parameters of spiking neurons were not included as trainable variables. By using the mollifier approximation [27], we developed a comprehensive framework that can tune both connectivity and spiking parameters using a gradient-descent method. Training spiking RNNs on multiple tasks using our method revealed functional specialization of excitatory and inhibitory neurons. More importantly, our approach allowed us to identify fast membrane voltage dynamics as an essential property required to encode incoming stimuli robustly for WM tasks.

Previous computational studies employing RNNs assumed that all the units in a network shared the same intrinsic parameters and optimized only synaptic connectivity patterns during training. Recent studies developed models that give rise to units with heterogeneous intrinsic properties. For example, a new activation function that is tunable for each neuron in a network was recently proposed [36]. In addition, we recently trained synaptic decay time constants in a rate RNN model [25]. Although these methods produce heterogeneous units, they do not incorporate parameters inherent to spiking mechanisms. Our method not only allows direct training of synaptic weights of spiking RNNs that abide by Dale’s principle, but also enables training of synaptic and intrinsic membrane parameters for each neuron.

Although our method was successful at training spiking RNNs with biological constraints, the gradient-based method employed in the present study is not biologically plausible. In cortical neural networks, local learning rules, such as spike-timing-dependent plasticity (STDP), were observed, but the gradient-descent algorithm used in our method is neither local to synapses nor local in time [16]. However, this non-locality allowed our method to train intrinsic membrane and connectivity parameters, creating biologically plausible neural architectures that solve specified problems. The learning algorithm for spiking neurons makes it possible to uncover neural dynamics hidden in experimental data [8, 29, 37], thus emphasizing that a biologically realistic model can be constructed by non-biological means.

Another limitation of our framework arises from our spiking neuron model. Although we were able to train models with heterogeneous neurons the leaky integrate-and-fire model used in the present study can only capture the dynamics of fast-firing neurons due to the lack of adaptation [38]. In particular, several other types of neurons, such as regular-firing and bursting neurons, are also common in cortical networks [39]. Applying our method to spiking neuron models with adaptation currents, such as those in Hodgkin-Huxley models model [40] and adaptive exponential integrate-and-fire model [41], will be an interesting next step to further investigate the role of neurons from various firing classes in information processing.

In summary, we provide a novel approach for directly training both connectivity and membrane parameters in spiking RNNs. Training connectivity and intrinsic membrane parameters revealed distinct populations only identifiable by their parameter values, thus enabling investigation of the roles played by specific populations in the computation processes. This lays the foundation for uncovering how neural circuits process information with discrete spikes and building more power-efficient spiking networks.

## Acknowledgements

We are grateful to Jorge Aldana for assistance with computing resources. This work was funded by the DARPA (W911NF1820259 to TJS) and National Institute of Mental Health (F30MH11560501A1 to R.K.). We also gratefully acknowledge the support of NVIDIA Corporation with the donation of the Quadro P6000 GPU used for this research. The funders had no role in study design, data collection and analysis, decision to publish, or preparation of the manuscript.

## Author contributions

Y.L., R.K., and T.J.S. designed the study and wrote the manuscript. Y.L. performed the analyses and simulations.

## Methods

### Spiking network structure and discretization

Our spiking RNN model consisted of *N* integrate- and-fire (LIF) units is governed by

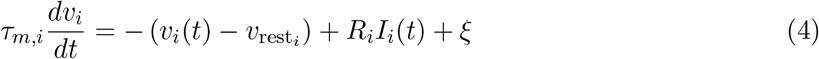

where *τ*_*m,i*_ is the membrane time constant of unit *i*, *v*_*i*_(*t*) is the membrane voltage of unit *i* at time *t*, *v*_*rest,i*_ is the resting potential of unit *i*, and *R*_*i*_ is the input resistance of unit *i*, and *ξ* is the membrane voltage spontaneous fluctuation. *I*_*i*_(*t*) represents the current input to unit *i* at time *t*, which is given by:

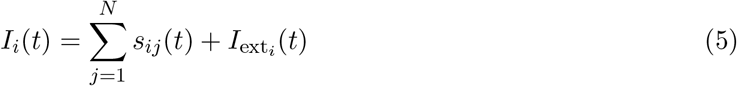

where *N* is the total number of units in the network, *s*_*ij*_(*t*) is the filtered spike train of unit *j* to unit *i* at time *t*, and *I*_ext_*i* (*t*) is the external current source into unit *i* at time *t*. For this study, *N* = 400 for all tasks and networks trained.

The external current ***I***_ext_(*t*) encodes the task-specific input at time *t*:

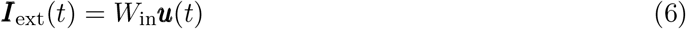

where the time-varying stimulus signals ***u***(*t*) ∈ ℝ^*N*_in_×1^ are fed into the network via *W*_in_ ∈ ℝ^*N* ×*N*_in_^, which can be viewed as presynaptic connections to the network that convert analog input into firing rates. *N*_in_ corresponds to the number of channels in the input signal.

We used a single exponential synaptic filter:

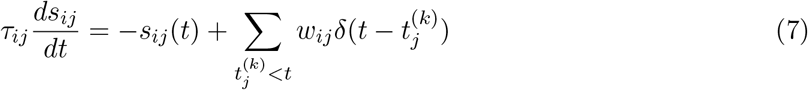

where *τ*_*ij*_ is the synaptic decay time constant from unit *j* to unit *i*, *w*_*ij*_ is the synaptic strength from unit *j* to unit *i*, 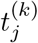 denotes the time of the *k*-th action potential of unit *j*, and *δ*(*x*) is the Dirac delta function. Once the membrane voltage of the unit *i* crosses its action potential threshold (*ϑ*_*i*_), its membrane voltage is brought back down to its reset voltage (*v*_*reset,i*_).

The output of our spiking model at time *t* is given by

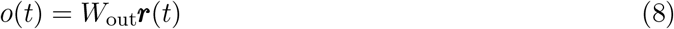

where *W*_out_ ∈ ℝ^1×*N*^ are the readout weights, and ***r***(*t*) ∈ ℝ^*N* ×1^, which can be interpreted as the firing rate of units, are given by

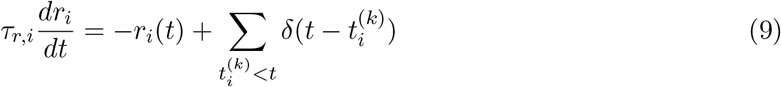

where *τ*_*r,i*_ is the synaptic decay time constant of firing rate estimate for unit *i*.

We converted the continuous-time differential equations to discrete-time iterative equations and used numerical integration (Euler’s method) to solve the equations. The membrane voltage ***v*** ∈ ℝ^1×*N*^ at step *n* + 1 is given by

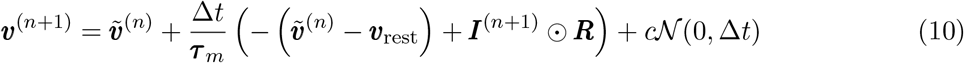

where Δ*t* is the sampling rate (or step size), which was set Δ*t* = 1 ms for this study, ***τ***_*m*_ ∈ ℝ^1×*N*^ is the membrane time constant, ***v***_rest_ ∈ ℝ^1×*N*^ is the resting potential, ʘ refers to Hadamard operation (element-wise multiplication), ÷ refers to the element-wise division, and ***R*** ∈ ℝ^1×*N*^ is the input resistance. The term 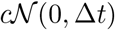 injects spontaneous membrane fluctuations, where 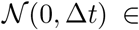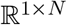 is a Gaussian random vector consisting of *N* independent Gaussian random variables with mean 0 and variance Δ*t*, and *c* is the scaling constant for the amplitude of fluctuations, set as *c* = 5 throughout the study.

There are two time-varying terms in Eq. 10, the membrane voltage after reset 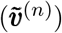 and input current (***I***^(*n*+1)^). The voltage reset in the LIF model after action potentials at step *n* is formulated as

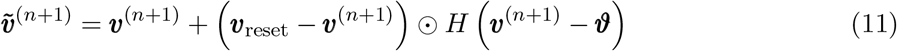

where ***v***_reset_ ∈ ℝ^1×*N*^ is the reset potential, ***ϑ*** ∈ ℝ^1×*N*^ is the action potential thresholds, and *H*(*x*) is the element-wise Heaviside step function. The term *H* (***v***^(*n*+1)^ − ***ϑ*** represents the spiking output activities at step *n* + 1. The input current at step *n* + 1 is given by

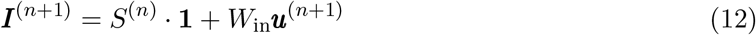

where **1**∈ ℝ^1×*N*^ is the column vector with all ones and *S*^(*n*)^ is the filtered spike train matrix at step *n*, which follows the iteration

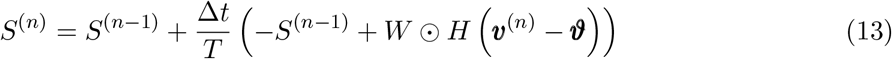

where *T* ∈ ℝ^*N* ×*N*^ is the matrix of synaptic decay time constants and *W* ∈ ℝ^*N* ×*N*^ is the matrix of synaptic strengths. Here, *W* ∈ ℝ^*N* ×*N*^ is a matrix and *H (**v***^(*n*)^ − ***ϑ***) ∈ ℝ^1×*N*^ is a row vector. The notation *A* ʘ ***v*** refers to element-wise multiplication of matrix *A* row by row with the row vector ***v***.

The output at step *n* + 1 is computed by

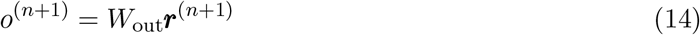

in which

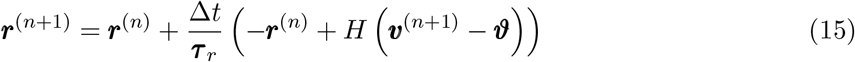

where ***τ***_*r*_ ∈ ℝ^1×*N*^ is the synaptic decay time constants of firing rate estimate.

### Training details

In this study, we only used the supervised backpropagation of errors learning algorithm. The loss function (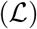) is defined in terms of the root mean square error (RMSE) with respect to a task-specific target signal (***z***) and the network output signal (***o***):

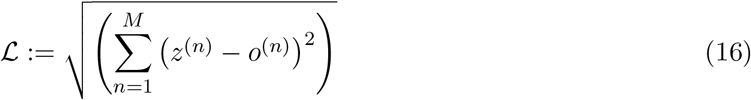

where *M* is the total time steps.

We used *Adaptive Moment Estimation (ADAM) stochastic gradient descent algorithm* [42] with mini-batch training. The mollifier gradient approximations were employed to address non-differentiability problem associated with the spiking process (see **Mollifier gradient approximations**). The learning rate was set to 0.01, the batch size was set to 10, and the first and second moment decay rates were 0.9 and 0.999, respectively. The trainable parameters include input weights (*W*_in_), synaptic strengths (*W*), readout weights (*W*_out_), synaptic decay time constants (*T*), membrane time constants (***τ*** _*m*_), input resistances (***R***), resting potentials (***v***_rest_), reset voltages (***v***_reset_), action potential thresholds (***ϑ***), and synaptic decay time constants for firing rate estimates (***τ*** _*r*_).

A *nonlinear projected gradient method* was used to constrain parameters within the biologically realistic ranges described in Supplementary Table 1. A linear projection map forces some solutions to be projected on the boundary. That is, there are always some units whose parameters take the min and max values of the constraint. On the other hand, a nonlinear projection guarantees that no values are on the boundary almost surely, a more realistic situation to consider. Specifically, to bound a parameter *p* at iteration *i* + 1 into the range [*p*_*min*_, *p*_*max*_], we have

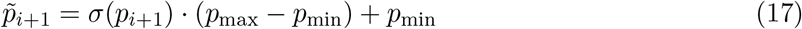

where 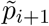 is the projected solution of parameter *p* at iteration *i* + 1, *p*_*i*+1_ is the unconstrained solution given by the gradient descent algorithm at iteration *i* + 1, *p*_max_ and *p*_min_ are the maximum and minimum values of parameter *p*, and *σ*(*x*) is the sigmoid function, defined as

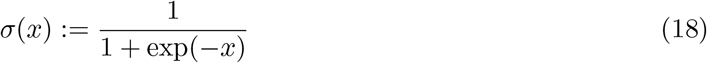

We *initialized all parameters*, except the input weights (*W*_in_), as samples from the standard Gaussian distribution with zero mean and unit variance, whereas the input weights were drawn from Gaussian distribution with zero mean and variance 400. This is because our input signals were bounded within the range [−1, 1], insufficient to bring the membrane voltage from the resting potential above the action potential threshold. Hence, to accelerate training, it was necessary to make sure units were excited by the input signals in the first place. The synaptic strength matrix (*W*) was also initialized sparse, with the percentage of connectivity being only 20%. We say the network successfully did the task if the output signal hits above +0.8 (or below −0.8) if the target output is +1 (or −1). We stopped training when the loss 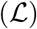 is less than 15 and the accuracy over 100 trials is above 95%.

The method proposed by Song et al. [29] was used to *impose Dale’s principle* with separate excitatory and inhibitory populations. The synaptic connectivity matrix (*W*) in the model was parametrized by

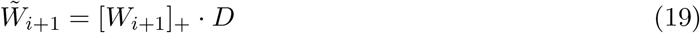

where 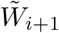 is the resulted matrix that encoded separate populations at update step *i* + 1, *W*_*i*+1_ is the solution given by the gradient descent algorithm at step *i* + 1, and [·]_+_ is the rectified linear unit (ReLU) operation applied at the end of each update step. The ReLU operation is to ensure that entries of the matrix are always non-negative before multiplied by the matrix *D*, as the negative weight connections update from gradient descent are pruned by the end of each update. The diagonal matrix (*D* ∈ ℝ^*N* ×*N*^) encode +1 for excitatory units and −1 for inhibitory units. The value of matrix (*D*) was randomly assigned before training according to a preset proportion between inhibitory and excitatory units, and the value *D* was fixed through the whole training process. The I/E units proportion in this study was 20% to 80%.

In order to capture the biologically realistic dynamics of SNNs, the *temporal resolution* (Δ*t*) was set to be no longer than the duration of absolute refractory period to ensure that the spiking activities are not affected by the numerical integration process. Therefore, we set Δ*t* = 1 ms during training. Due to the vanishing gradient problem occurring in training RNNs [43], with Δ = 1 ms, it is impossible to train tasks with duration longer than 1 second (i.e., *M* > 1000). It is notable that in the above formulation, only membrane time constant (*τ*_*m*_) and synaptic time decay (*τ*) are dependent on the sampling rate (Δ*t*; Eq. 10 and Eq. 13). Hence, after the models are trained, we can make sampling rate (Δ*t*) smaller (i.e., having finer temporal resolution) while still keeping the same dynamics of the trained networks. Increasing Δ*t* by a factor is equivalent to decreasing *τ* and *τ*_*m*_ altogether by the same factor, as *τ* and *τ*_*m*_ are inversely proportional to Δ*t* in Eq. 10 and Eq. 13. Hence, to train a network performing tasks with duration longer than 1 second, we need to make the temporal resolution coarser (i.e., increasing Δ*t* by a factor *s*) so that with the same trainable range of time steps (i.e., a fixed *M* ≤ 1000), the duration of task becomes longer by the same factor *s*. This “decrease in temporal resolution” can be interpreted as shortening *τ* and *τ*_*m*_ instead of an actual decrease in temporal resolution. Applying this trick enables us to train tasks with arbitrary duration by re-scaling the ranges of *τ* and *τ*_*m*_ into a smaller one while still making the spiking activities biologically realistic. In practice, we simply scaled down *τ* and *τ*_*m*_ by a factor *s* = 3 with a fixed number of time steps (*M*), and later during the testing stage, we re-scaled *M, τ* and *τ*_*m*_ up by the same factor *s*.

### Mollifier gradient approximations

In the above formulation, the Heaviside step function *H*(*x*) is not continuous. As a result, the loss function 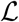 is not differentiable. This poses the major problem when applying the traditional backpropagation algorithm for training neural networks, because the backpropagation algorithm uses gradient descent methods that require the function being minimized to be differentiable, or at least to be continuous. However, the derivative of Heaviside step function *H*(*x*) is Dirac Delta function *δ*(*x*), which is 0 everywhere except at 0, where the function value is ∞. It is difficult to use this derivative for the gradient descent methods because the value of the gradients is 0 almost everywhere.

To address the discontinuity problem, we employed mollifier gradient method proposed by Ermoliev et al. [27]. The method can be applied to any strongly lower semicontinuous functions to find local minima following an iterative gradient descent in which the gradients change over iterations based on averaged functions derived from the original objective function. The family of averaged functions *f*_*ε*_ of function *f* is defined by convolution of *f* with a mollifier: *ψ*_*ε*_

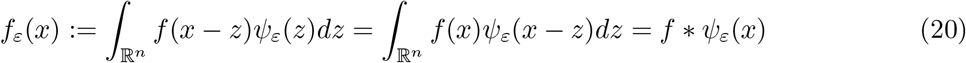

where *ψ*_*ε*_ ∈ {*ψ*_*ε*_ : ℝ^*n*^ → ℝ_+_, *ε* > 0}, a family of compactly supported (generalized) functions named *mollifiers* that satisfy

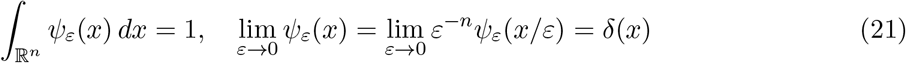

It was shown that for any strongly lower semicontinuous functions *f*, the averaged functions *f*_*ε*_ epi-converge to *f* as *ε* → 0, a type of convergence that preserves the local minima and minimizers. Therefore, it is possible to use the gradients of averaged functions to minimize the original lower semicontinuous functions and find the local minima. We used the conventional family of mollifiers obtained by normalizing a probability density function *ψ*:

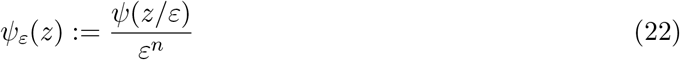

In our case, *n* = 1 as the domain of *H*(*x*) is the real line:

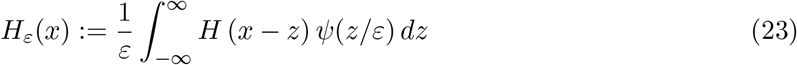

For any *ε* > 0, the gradient of *H*_*ε*_ (*g*(*x*)) with respect to parameter *p* is given by

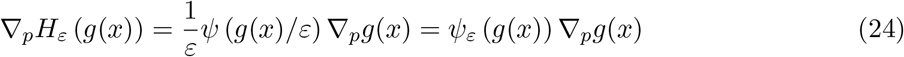

where *ψ* is some symmetric density function and *g*(*x*) is any function with R as its codomain. Since our goal was not to find a local minimum *x*\* that satisfies the optimality condition 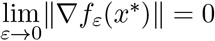 as defined by Ermoliev et al., but rather to minimize the loss function for its value to be sufficiently small so that the network can perform the task correctly, we did not vary the gradients during the minimization process. Instead, we fixed an approximation of the gradient and used the approximation throughout the training process. We chose the normalized box function, i.e., the density function of uniform distribution 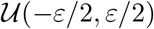, as the kernel,

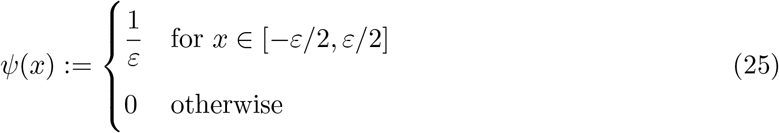

and fixed *ε* = 5.

We found no difference in the trained models with different choices of *ε*, as long as the value was large enough to keep the gradients active so that gradients did not vanish through time steps. There was also no difference between models trained with fixed *ε* and those trained with the original scheme in Ermoliev et al. where *ε* → 0 as the number of iterations increases. The purpose for fixing the value of *ε* was to compare the training epochs (iterations) among the retraining paradigms (see Fig. 3) with the same gradient.

### Re-training models for DMS task

To test whether intrinsic properties optimized for one WM task are generalizable to other tasks that also require WM, we re-trained our models to perform the DMS task with all intrinsic properties fixed. In contrast to the training paradigm described in the previous sections, the trainable parameters for re-training only include input weights (*W*_in_), synaptic strengths (*W*), and readout weights (*W*_out_). Each of the 20 RNNs trained for each of the four tasks (non-WM integration, delayed integration, DMS, and DIS tasks) used in this study was re-trained to perform the DMS task.

To test whether synaptic decay time constants (*τ*) and membrane time constants (*τ*_*m*_) are the most crucial parameters for transferability of WM tasks, we repeated the re-training procedure with both *τ*_*m*_ and *τ* either fixed or optimized for the non-WM RNNs. The RNNs optimized to perform the context-based input integration task were used for re-training under two schemes: the tuned scheme and the frozen scheme. For the tuned scheme, the trainable parameters include input weights (*W*_in_), synaptic strengths (*W*), readout weights (*W*_*out*_), synaptic decay time constants (*T*), membrane time constants (***τ*** _*m*_), and synaptic decay time constants for firing rate estimates (***τ*** _r_). For the frozen scheme, the trainable parameters include input weights (*W*_in_), synaptic strengths (*W*), readout weights (*W*_out_), input resistances (***R***), resting potentials (***v***_rest_), reset voltages (***v***_reset_), and action potential thresholds (***ϑ***).

### Units function analysis

For Fig. 4, we manipulated *τ*_*m*_ and *τ* during different epochs of the DMS task paradigm to investigate if fast membrane and slow synaptic dynamics are responsible for WM maintenance. For each of the RNNs trained from the DMS task, we first divided the population into two subgroups based on their *τ*_*m*_ values. The short *τ*_*m*_ group contained units whose *τ*_*m*_ was smaller than the median value of *τ*_*m*_ of all units in the RNN, while the long *τ*_*m*_ group contained units whose *τ*_*m*_ was greater than the median value. The average median value of *τ*_*m*_ across all 20 models was 19.64 ± 2.45 ms. During each of the four epochs (fixation, first stimulus, delay, and second stimulus), we inhibited the two *τ*_*m*_ subgroups separately by hyperpolarizing them and then assessed the task performance. The hyperpolarization was done by setting the membrane voltage *v* = −100 mV for the intended subgroup of units. Similar to the training stage, we say that the network successfully did the task if the output signal hits above +0.8 (or below −0.8) if the target output is +1 (or −1). If the target output is between −0.8 and +0.8, the network is considered having no response. If the output signal is above +0.8 (or below −0.8) while the target output is −1 (or +1), we say that the network gives an incorrect response.

We conducted a similar analysis based on two subgroups of synapses derived from a quartile split of synaptic decay time constant (*τ*). The short *τ* group contained synapses whose *τ* was smaller than the 25th percentile of all *τ* in the RNN, while the long *τ* group contained synapses whose *τ* was greater than the 75th percentile. The average 25th percentile across all 20 models was 25.36 ± 2.40 ms, and the average 75th percentile was 66.18 ± 1.17 ms. The targeted subgroup of synapses was suppressed by setting the connection strength *w* = 0 during each of the four epochs of DMS task.

## Code availability

The implementation of our framework and the codes to generate all the figures in this work are available at https://github.com/y-inghao-li/SRNN/

## Data availability

The trained models used in the present study are available as MATLAB-formatted data at https://github.com/y-inghao-li/SRNN/

## Supplementary Table

**Supplementary Table 1:**
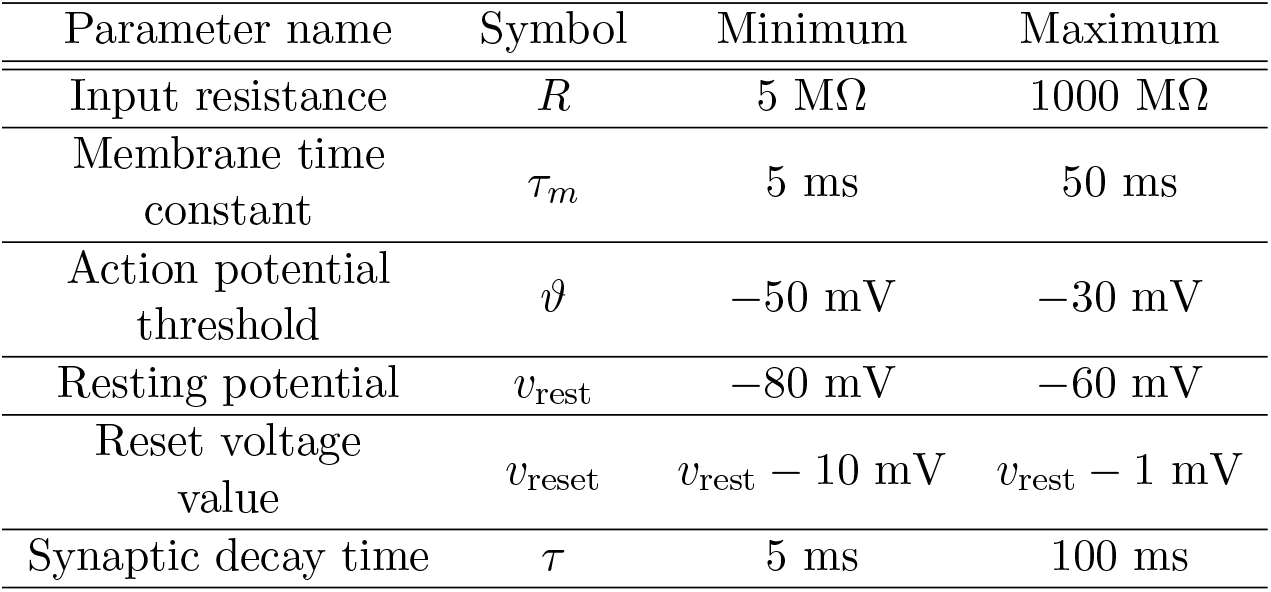
Parameter values used for this study. To keep the constraint *v*_rest_ > *v*_reset_, we trained the afterhyperpolarization (AHP) potential with range from −10 mV to −1 mV, so the value of *v*_reset_ is dependent upon the value of *v*_rest_.

